# Simple and rapid lectin-based detection of native glycoRNAs

**DOI:** 10.1101/2024.12.09.627596

**Authors:** Yong Li, Yisong Qian, Tianhua Lei, Paula Monaghan-Nichols, Mingui Fu

**Affiliations:** Department of Biomedical Science, School of Medicine, University of Missouri Kansas City, Kansas City, MO 64108, USA; Department of Anesthesia, The first affiliated hospital, Nanchang University, Nanchang, Jiangxi 332306, China; Key Lab for Arteriosclerology of Hunan Province, International Joint Laboratory for Arteriosclerotic Disease Research of Hunan Province, Institute of Cardiovascular Disease, University of South China, Hengyang 421001, China; Shock/Trauma Research Center, School of Medicine, University of Missouri Kansas City, Kansas City, MO 64108, USA

**Author notes:** Corresponding author: Mingui Fu.

## Abstract

The current detection methods for glycoRNAs include metabolic labeling of living cells or animals and rPAL that directly detects native glycoRNAs by periodate oxidation and aldehyde ligation. Both of them have some limitations. Here we reported a simple and rapid detection of native glycoRNAs by using lectins. The method involves in several simple procedures: isolation of total RNA, northern blotting and detection with lectins. The advantages of this method include high sensitivity, very simple procedures and broader applications. We used this method detecting glycoRNA expression in the RNA samples from different species. We also detect the glycoRNA expression across the varieties of mouse and human tissues and compared with other detection methods. We are in the first to detect free glycoRNAs in the varieties of human biofluids. Overall, this simple and rapid method will provide a new tool for study of glycoRNA biology and clinical diagnosis in future.

---

It was recently reported that RNAs can be glycosylated, and such glycosylated RNAs (called glycoRNAs) are located on the outer cell surfaces including human stem cells and several cancer cell lines (1). Zhang et al. further demonstrated that the cell surface glycoRNAs are important molecules that mediate neutrophils recruitment into inflammatory area by selective interaction with adhesion molecules on vascular endothelial cells such as P-selectin (2). We recently found that human monocytes expressed robust glycoRNAs and it mediates the interaction of human monocytes and endothelial cells via binding to siglec-5 (3). Although there are still many things unclear about the newly discovered cell surface RNAs, evidence suggests that there are glycan-modified small noncoding RNAs, which are synthesized inside cells and transported to cell plasma membrane. These intriguing discoveries open a new avenue for studying the molecular mechanisms that control immune response, cell-cell interaction, pathogen-host interaction, embryogenesis and cancer cell metastasis (4-6).

The current detection methods for glycoRNAs include metabolic labeling of living cells or animals (1, 2) and rPAL that directly detects native glycoRNAs by periodate oxidation and aldehyde ligation (7). Both of them have some limitations. For example, metabolic labeling is dependent on living cells or animals that take up the labeling molecule before their biosynthetic processing and incorporation into nascent biopolymers, like glycans. This limits metabolic labeling method only can use in contexts where access to organism or cells with a reporter is possible. rPAL method resolved this problem that can be used in any samples containing glycoRNAs. However, rPAL only detects sialoglycoRNAs, but not the other glycoforms. The labeling and purification steps make it not suitable for detection of human samples that contain less RNAs such as serum and other biological fluids.

GlycoRNA is a hybrid molecule containing glycan-modified small non-coding RNA. The principle for detection of glycoRNAs is “*purifying RNAs + detecting glycan = detection of glycoRNAs*”. When total RNAs are isolated from cells or tissues, the other molecules including proteins, DNA and lipids have been removed. The total RNA was separated by denaturing formaldehyde gel electrophoresis and then blotting on membrane. Then we need to detect glycan to determine the exist of glycoRNAs. In the world, there are many proteins that specifically bind to glycan chains, named lectins (8). To investigate which lectin that is suitable for detecting glycoRNAs, we purchased 20 different lectins from Vector Laboratories. The total RNAs isolated from THP1, HeLa and HEK293 cells were used for screening these lectins. Most of lectins tested can detect glycoRNAs (Extended data Fig.1a). However, WGA, LEL, DSL and STL give more strong bindings. Then, we compared the binding activity of WGA, LEL, DSL and STL, which showed that LEL is the strongest one in binding to glycoRNAs (Extended data Fig. 1b). LEL, lyopersicon esculentum lectin, is an agglutinin from tomato, which binds to the sugars containing N-acetylglucosamine (Extended data Fig. 2a). The protein LEL has 339 amino acids containing four chitin-binding domains (Extended data Fig. 2b). Normally, lectin crystal structure showed four specific domains for binding with glycans (Extended data Fig. 2c).

Next, we optimized the experimental conditions for LEL-based detection of glycoRNAs. The total RNAs isolated from THP1 cells was used for all the optimizations. First, we screened the blocking buffers. Protein-free blocking buffer purchased from Li-Cor company generated more cleaner background compared to 0.5% BSA and 5% milk. 0.5% BSA can also be used, however, general skim milk cannot be used as it gives dark background due to containing other sugars (Extended data Fig. 3a). Two types of membranes (nitrocellulose membrane and nylon membrane) were tested. Nylon membrane that is usually good for northern blot cannot be used in this detection as it generated dark background (Extended data Fig. 3b). Three types of transfer buffers were also tested including one from Invitrogen, 2XSSC, and 3M NaCI (pH 1.0). Though the RNA transfer efficiency is highest using 3M NaCI as described by Xie et al. (7), but the transfer buffer from Invitrogen gave cleaner background. 2XSSC can also be used, as it gave good background, and it is the cheapest one (Extended data Fig. 3c). Two types of agaroses (agarose-LE from Invitrogen and general agarose from Themo-Fishers) were tested. The general agarose cannot be used for LEL but can be used for metabolic labeling method. Ultra-pure agarose-LE must be used for LED (Extended data Fig. 3d). Next, we optimized the concentration of LEL and incubation times. 3 μg/ml of LEL is good for generating strong signal (Extended data Fig. 3e). Though 24 hours of LEL incubation generated strongest signal, 2-4 hours of incubation also generated applicable signals (Extended data Fig. 3f).

To verify the specificity of lectin-based detection (LBD), total RNAs were isolated from THP1 cells treated with or without NIG-1, an inhibitor of oligosaccharyltransferase, or Kifunensine, an inhibitor of α-mannosidase-I. Previous report showed that both inhibitors repressed the synthesis of glycoRNAs (1). In the two cases we saw loss of glycoRNA-signal in THP1 that were treated with these inhibitors (Extended data Fig. 3g). Next, we compared the sensitivity of LBD with metabolic labeling and rPAL. The minimum RNA that can be detectable by metabolic labeling and rPAL is 0.5 μg; whereas the minimum RNA that can be detectable by LBD is 0.1 μg (Fig. 1a), suggesting that LBD is the most sensitive method. As we recently reported (3), there are two glycoforms of glycoRNAs can be detected in THP1 cells by metabolic labeling. However, the small form of glycoRNAs only appeared in metabolic labeling detection. GlycoRNA-S in THP1 cells cannot be detected by LBD and rPAL. One possible reason is that lectin may be not able to bind the small glycan chain of glycoRNA-S. The other possibility is that glycoRNA-S is generated by metabolic probe. Nevertheless, the identity of the small glycoform observed in THP1 cells need to be further clarified.

**Figure 1.**
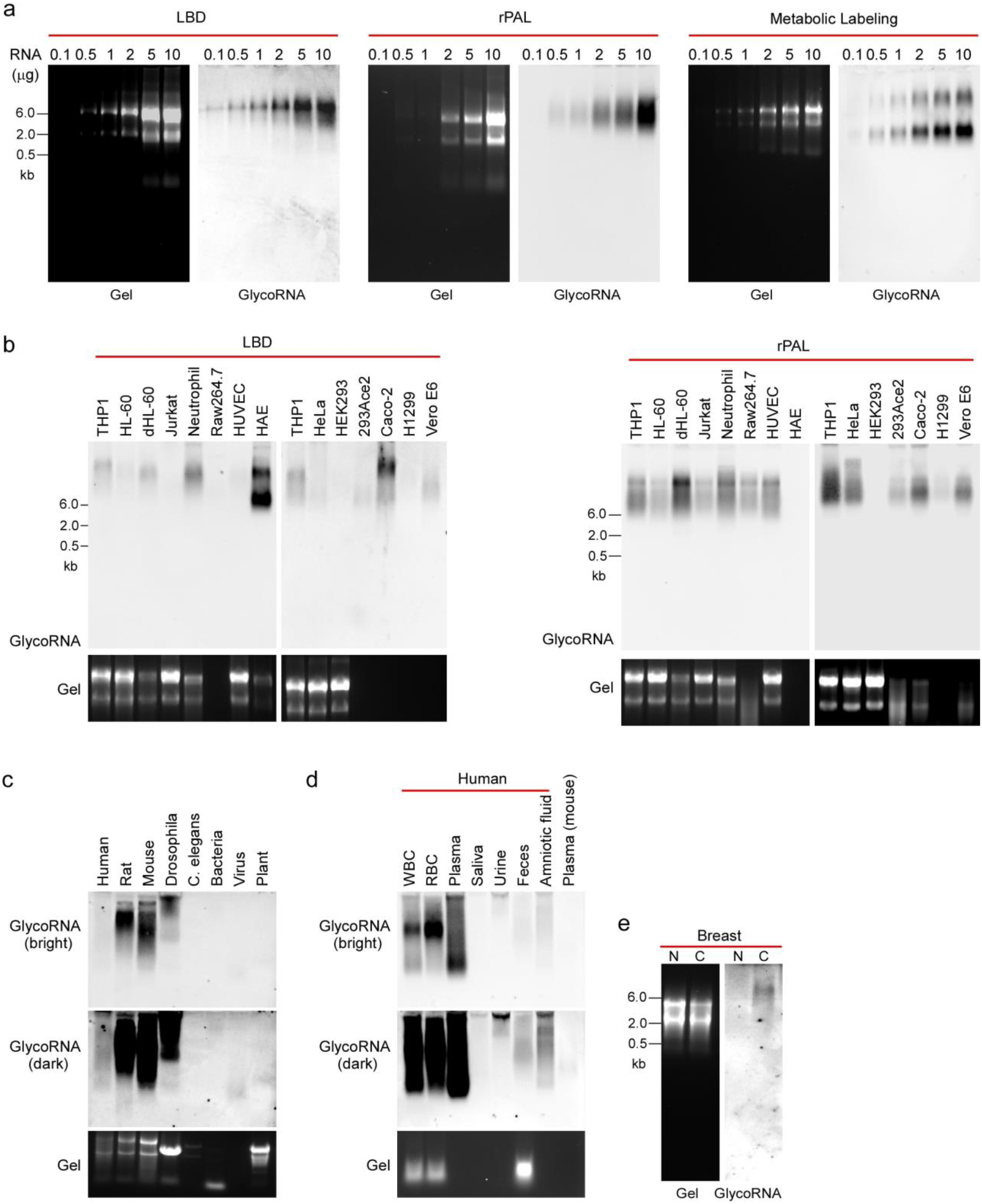
Development of lectin-based detection of native glycoRNAs and its application. (a) The sensitivity of three methods as indicated were compared. The RNAs were from Ac4ManNAz-labeled or unlabeled THP1 cells. (b) The expression of native glycoRNAs in the cell lines as indicated was detected by LBD and rPAL. 10 μg total RNAs were used. (c) 2-10 μg total RNAs from different species as indicated were analyzed by LBD. (d) 2-5 μg RNAs isolated from human blood cells, plasma, saliva, urine, feces, amniotic fluid and mouse plasma were analyzed by LBD. (e) The total RNAs from human normal breast tissue (N) or breast cancer tissue (C) were analyzed by LBD.

The step-by-step protocol for lectin-based detection of native glycoRNAs was finalized (Extended data Fig. 4). Next, we tested if this method can be used for detection of native glycoRNAs from different cell lines. Total RNAs were isolated from different cell lines as indicated in Fig. 1b. The glycoRNA expressed was higher in HAE, Caco-2, THP1, dHL-60, mouse neutrophils, HeLa and Vero E6 cells, lower in HL-60, Jurkat, Raw264.7, HUVECs, H1299 and HEK293 cells (Fig. 1b). These results are consistent with that by rPAL. Interestingly, though some RNA samples showed heavy degradation (RAW264.7, 293Ace2, Caco-2, H1299 and Vero E6), the detection for glycoRNA expression is not affected, suggesting that glycoRNAs are not sensitive to RNase-mediated degradation. One possible reason is that glycoRNAs may be protected by its self-glycan chains. Another possible reason is that glycoRNAs may be protected by some RNA-binding proteins (2, 9). Take together, both LED and rPAL generated very consistent results.

As LEL is purified from plant, theoretically it should be able to bind the glycan chains from different species. Total RNAs isolated from human (thymus), rat (thymus), mouse (thymus), Drosophila (whole), C. elegans (whole), bacteria (E. coli, DH5α), virus (SARS2, XBB1.5) and plant (Arabidopsis) were used for detection. Both rat and mouse thymus expressed robust glycoRNAs as described previously (3), however, human thymus displayed weak signals. Interestingly, we detected strong signal from Drosophila, and very weak signals from virus and plant when overexposure (Fig. 1c). This result suggest that we are in the first to detect the glycoRNAs in a non-mammalian animal (Drosophila).

As glycoRNAs are localized on the outer surface of cells, theoretically they may be released into blood by RNase degradation. To investigate whether there are free glycoRNAs existed in plasma and other biofluids, we collected RNA samples from human plasma, saliva, urine, feces, amniotic fluid as well as WBC and RBC and mouse plasma. As shown in Fig. 1d, there is robust glycoRNA expression in human white blood cells as described previously (7). Surprisingly, we also detected robust glycoRNAs from red blood cells and plasma. Weak signals were also observed from urine, feces and amniotic fluid. These results suggest that there are free glycoRNAs existed in human plasma and other biofluids, which may be able to serve as a biomarker for diagnosis of huma diseases.

Next, we tested if LBD can also be used for RNA samples from pathological settings. We purchased total RNAs of human breast and breast cancer from Biochain. There is no glycoRNAs expression in normal human breast tissues, however, there is detectable glycoRNAs expression in human breast cancers (Fig. 1e). Taken together, this method can be used to detect native glycoRNAs from both human and mouse tissues under physiological and pathological conditions.

To investigate the expression patterns of glycoRNA across mouse and human tissues, we fist isolated total RNAs from 22 tissues from adult C57/BL6 mice. The native glycoRNAs expressed in these tissues were detected by LED and rPAL. As shown in Fig. 2a, glycoRNAs are highly expressed in immune organs including thymus, spleen, lymph nodes, bone marrow and WBC; guts including esophagus, stomach, intestine and colon; and other organs including brain, heart, WAT, RBC etc. GlycoRNAs were not detectable in cerebellum, lung, liver, kidney, eye, BAT, muscle and skin. Both LED and rPAL generated very consistent results. To compare with metabolic labeling, adult C57/BL6 mice were intraperitoneally injected with 300 mg/kg/day of Ac4ManNAz for 2 days as described by Flynn et al (1). Total RNA was isolated from the tissues as indicated and analyzed by the procedures as described by Flynn et al. The expression patterns of glycoRNAs in mouse tissues generated by metabolic labeling were similar to that generated by LED and rPAL described above. However, glycoRNAs were not detectable in brain and bone marrow by metabolic labeling. We postulate that the difference may be due to that the metabolic probe (Ac4ManNAz) cannot pass the blood-brain barrier or blood-bone marrow barrier. In addition, we observed that the results from metabolic labeling are very unstable, which indicates that metabolic labeling may be affected by many factors. The quantification of glycoRNA levels in each tissue were presented and compared (Extended data Fig. 6a&c). Overall, the results from LED and rPAL are very comparable, the results from metabolic labeling are similar but have some differences due to its limitations. Interestingly, the glycoRNAs are highly expressed in mouse cerebral cortex, midbrain, four colliculi, hippocampus, hypothalamus, striatum, and spinal cord, but not expressed in cerebellum (Extended data Fig. 5a&b). To profile the expression patterns of glycoRNAs in human tissues, we purchased 20 RNA samples from different tissues commercially, the other three RNA samples from WBC, RBC and placenta were isolated by my lab. First, we used LBD to profile the glycoRNAs in these tissues. Surprisingly, though the expression of glycoRNAs from WBC, RBC, placenta, brain, thymus, artery were also detected in human, the expression of glycoRNAs from other tissues such as guts, heart, spleen, lymph node, bone marrow and adipose were not detected (Fig. 2b). As the highly expressed tissue RNAs such as WBC, RBC and placenta were freshly isolated by my lab, all commercial derived RNAs showed weak or no signal, we postulate that the RNA isolation procedures for human tissues may be a factor to affect the detection. We also used rPAL to detect the glycoRNAs in some human tissues. The expression of glycoRNAs in human thymus, lung, and placenta is consistent with the results generated by LBD. However, rPAL also detected signals from colon, bladder, intestine, stomach and esophagus, confirming that glycoRNAs are also expressed in human guts (Extended data Fig. 6b). We also isolated some tissue RNAs from rat and the expression patterns of glycoRNAs were similar with that in mice (extended data Fig.5c. We also used WGA, a lectin from wheat, to detect the expression of glycoRNAs in some mouse tissues, which is similar with the results generated by LEL (extended data Fig. 5d). Finally, we treated total RNAs from mouse colon with or without proteinase K or RNase. Proteinase treatment didn’t change the levels of glycoRNAs, however, RNase treatment resulted disappear of glycoRNA signal (extended data Fig. 5e). Together, the predominant expression of glycoRNAs in immune organs, guts, brain, heart, aorta and adipose suggests that glycoRNAs may be critical in the regulation of immune response, pathogen-host interaction, gut barriers, central nerve system, cardiovascular and adipose functions.

**Figure 2.**
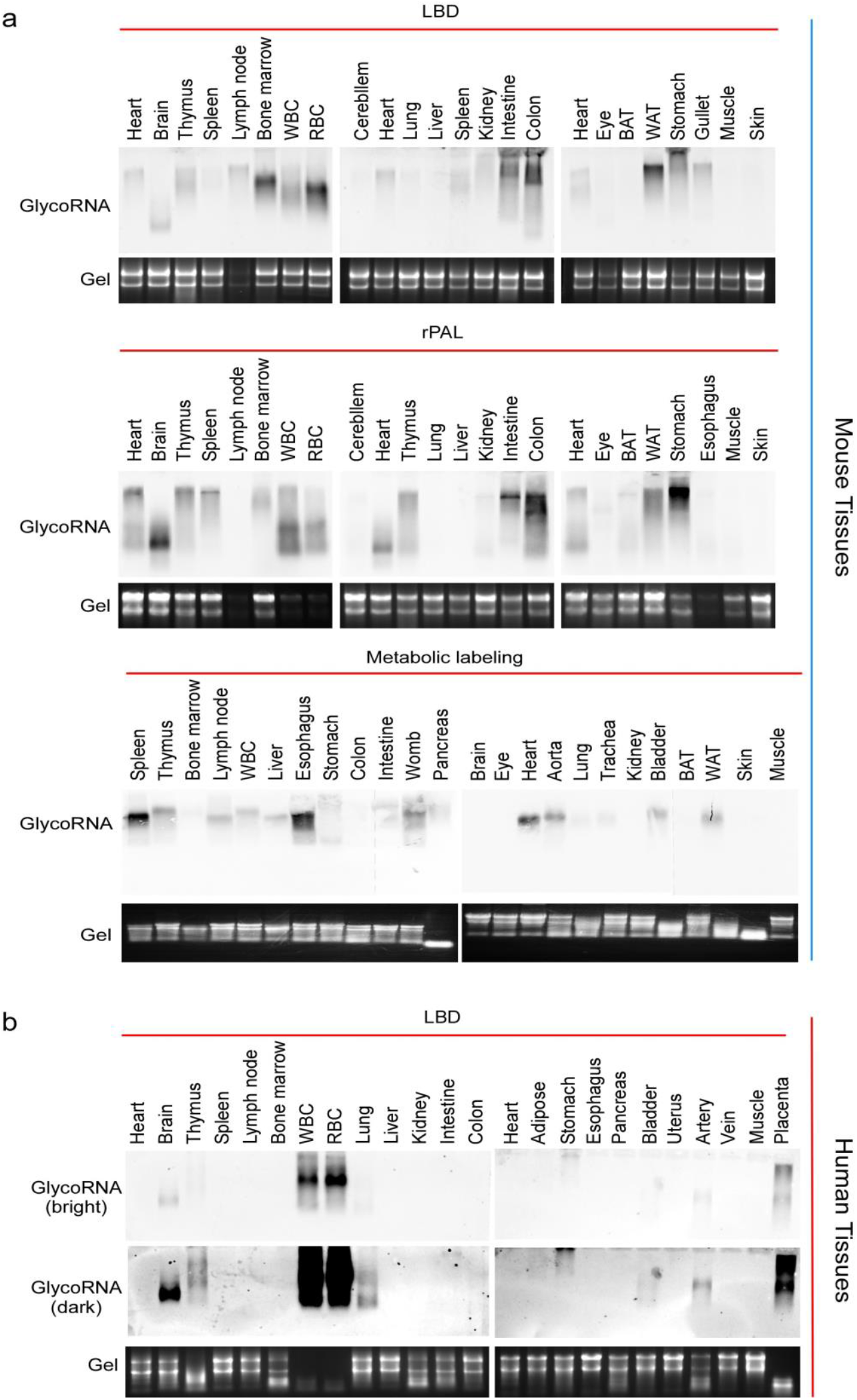
Expression patterns of glycoRNAs in mouse and human tissues. (a) The expression pattern of glycoRNAs in mouse tissues were analyzed by three methods: LBD, rPAL and metabolic labeling as indicated. 10-15 μg total RNAs were used. The experiments were repeated at least twice. (b) The expression pattern of glycoRNAs in human tissues were analyzed by LBD. 8-10 μg total RNAs were used.

In summary, metabolic labeling is based on inserting a chemical modified monosaccharide into glycan chains in living cells or animals (extended data Fig.7). After purifying RNA and separating by denaturing gel, the glycoRNAs are measured by detecting the modified sialic acids. rPAL is chemically modifying the atom groups on sialic acid in vitro after purifying RNAs and then purifying again and detecting the modified sialic acids. LBD is directly detecting the glycoRNAs after RNA purifying and separating. Compared with the metabolic labelling and rPAL, the LBD method has several advantages: 1) higher sensitivity: after optimizing the experimental conditions. LBD showed more higher sensitivity than metabolic labelling and rPAL. The minimum amount of RNA that can be detectable by LBD is 100 ng, which is better than metabolic labeling and rPAL (500 ng in minimum). 2) broader application: LBD potentially can detect different glycoforms of glycoRNAs, whereas metabolic labelling and rPAL only can detect sialoglycoRNAs. Thus, LBD could detect the glycoRNAs from any species such as Drosophila, C. elegans, bacteria, viruses or plants. 3) More simple procedures and easier to perform: LBD avoids labeling and purification steps. 4) In addition, we observed that there was abundant glycoRNAs expression in mouse brain detected by LBD, however, the glycoRNAs in mouse brain is undetectable by metabolic labeling, which may be due to the labeled metabolites cannot pass the blood-brain barrier. Thus, the LBD is not affected by the metabolic factors. We also noted that there are some limitations with LBD. 1) As lectins do not bind to monosaccharide or simple glycoforms (<3 sugar molecules, ref. 10), this method cannot detect monosaccharide-conjugated RNAs; 2) The method cannot detect the glycoRNAs from fixed tissue slides or cell membranes of living cells; 3) The method is not quantitative (may be semi-quantitative). There is a report to image glycoRNAs on single cell (11). However, it needs specifically synthesized probe. We see opportunities to directly visualize glycoRNAs with imaging techniques by modifying the lectin probes. In the future, we may develop a lectin-based ELISA method to directly detect the glycoRNAs from human plasma, serum, urine and other biological fluids.

## METHODS

### Animals

C57/BL6 mice and SD rats were used in this study. The animals were euthanized by standard procedures and the tissues were quickly harvested within 2 mins and put into liquid nitrogen to frozen and then transported to −80°C for store. The protocols were approved by UMKC IACUC.

### Cell culture

All cells were grown at 37°C and 5% CO_2_. HeLa, HAE, HEK293, 293ACE2, Caco-2, H1299, vero E6 and Raw264.7 cells were cultured in DMEM media supplemented with 10% fetal bovine serum (FBS) and 1% penicillin/streptomycin (P/S). THP1, HL-60, Jurkat cells were cultured in RPMI-1640 media with 1 mM HEPES, 1 mM Sodium Pyruvate, 0.001% β-ME and glutamine supplemented with 10% FBS. dHL-60 is obtained by incubation of HL-60 with 3% DMSO for 3 days. HUVECs were cultured in EBM2 growth media (Lonza).

### Metabolic labeling of the cells

Stocks of N-azidoacetylmannosamine-tetraacylated (Ac_4_ManNAz, Tocris Bioscience) were made to 500 mM in sterile dimethyl sulfoxide (DMSO). In cell experiments ManNAz was used at a final concentration of 100 μM. Working stocks of glycan-biosynthesis inhibitors were all made in DMSO at the following concentrations and stored at −80°C: 10 mM NGI-1 (Sigma), 10 mM Kifunensine (Kif, Sigma), All compounds were used on cells for 24 h and added simultaneously with ManNAz for labeling.

### Metabolic labeling in mouse models

All experiments were performed according to guidelines established by the University of South China IACUC committee. C57BL/6 mice were crossed and bred in house. ManNAz was prepared by dissolving 100 mg ManNAz in 830 μL 70% DMSO in phosphate buffered saline (PBS), warming to 37°C for 5 min, and then sterile filtering using 0.22 μm Ultra free MC Centrifugal Filter units (Fisher Scientific); this solution was stored at −20°C. Male C57BL/6 mice (8-12 weeks old) were injected once-daily, intraperitoneally with 100 μL of ManNAz (dosed to 300 mg ManNAz/kg/day), while control mice received the vehicle alone. After 2 days, mice were euthanized, and their tissues were harvested. The organs were pressed through a nylon cell strainer and resuspended with PBS to create a single cell suspension. RNA was collected as described below.

### RNA extraction and purification strategies

For total RNA isolation, TRIzol reagent (Thermo Fisher Scientific) was always used as a first step to lyse and denature cells or tissues. After homogenization in TRIzol by pipetting, samples were incubated at 37°C for 10 mins to further denature non-covalent interactions. Phase separation was initiated by adding 0.2× volumes of 100% chloroform, vertexing to mix well, and finally spinning down at 12,000x g for 15 min at 4°C. The aqueous phase was carefully removed, transferred to a fresh tube and mixed with equal volume of isopropanol. The mixture was put in 4°C for 10 mins and then centrifuged at 12,000x g for 10 min at 4°C. The RNA pallet was washed by 1 ml of 75% ethanol (EtOH). The RNA pellet was dry at room temperature and dissolved by RNase-free water. After enzymatic treatment or biotin-conjugating, the samples were always purified by Zymo column. For isolation of RNA from plasma and biofluids, 250 μl of plasma or biofluids were mixed with 750 μl TRIzol-LS (Invitrogen) and pipette several times. The following procedures are same as described above. For isolation of urine RNAs, the urine was concentrated by Amicon filter sets (MM>9000 da) by 5 folds and then performed the same procedures as the other biofluids. The isolation of feces RNAs, the feces were mixed with equal volume of RNase-free water and shake for 15 mins. After centrifuge for 10 mins, the supernatant was subjected to the same procedures as above.

### Enzymatic treatment of RNA samples

To digest RNA the following was used: 1 μL of RNase cocktail (0.5U/μL RNaseA and 20U/μL RNase T1, Thermo Fisher Scientific) was added into 20 μl of total RNA at 37 °C for 10 mins. The reaction mixture was purified by Zymo column. To treat RNA by proteinase K, 1 μL of proteinase K (20 mg/ml) was added into 20 μl of RNA and incubated at 37 °C for 45 mins. The reaction mixture was purified by Zymo column.

### Copper-free click conjugation to RNA

Copper-free conditions were used in all experiments to avoid copper in solution during the conjugate of biotin to the azido sugars. All experiments used dibenzocyclooctyne-PEG4-biotin (DBCO-biotin, Sigma) as the alkyne half of the cycloaddition. To perform the SPAAC, RNA in pure water was mixed with 1x volumes of “dye-free” Gel Loading Buffer II (df-GLBII, 95% Formamide, 18 mM EDTA, and 0.025% SDS) and 500 μM DBCO-biotin. Typically, these reactions were 30 μL df-GLBII, 27 μL RNA, 3 μL 10 mM stock of the DBCO reagent. Samples were conjugated at 55°C for 10 min to denature the RNA and any other possible contaminants. Reactions were stopped by adding 2x volumes (120 μL) of RNA Binding Buffer (Zymo), vertexing, and finally adding 3x volumes (180 μL) of 100% EtOH and vortexing. This binding reaction was purified over the Zymo column as instructed by the manufacturer and analyzed by gel electrophoresis as described below.

### RNA gel electrophoresis, blotting, and imaging

Blotting analysis of ManNAz-labeled RNA was performed according to the procedures described by Flynn et al. (1) with the following modifications. RNA on the column was eluted by 18 μl H_2_O and then add 18 μl of df-GLBII with EB (Thermo Fisher Scientific). To denature, RNA was incubated at 55°C for 10 min and crashed on ice for 3 min. Samples were then loaded into a 1% agarose-formaldehyde denaturing gel (Northern Max Kit, Thermo Fisher Scientific) and electrophoresed at 60 V for 60 min. Total RNA was then visualized in the gel using iBright 1500 (Invitrogen). RNA transfer to NC membrane (0.45 μm) occurred as per the Northern Max protocol for 2-16 h at 25°C. After transfer, RNA was crosslinked to the NC using UV-C light (0.18 J/cm^2^). NC membranes were then blocked with Protein Free Blocking Buffer, PBS (Li-Cor Biosciences) for 60 min at 25°C. After blocking, the blot was incubated with anti-Biotin-HRP (1:1000) for 2-16 h at 4°C. Excess anti-biotin-HRP was briefly washed from the membranes by 0.1% Tween-20 (Sigma) in 1x PBS. NC membranes were imaged on iBright 1500 (Invitrogen).

### rPAL

rPAL was performed as described by Xie et al (7). Briefly, 14 μl blocking buffer (1μL 32 mM mPEG3-Ald (Creative labs), 7 μL 2M MgSO_4_ and 6 μL 2 M NH_4_OAc pH5 (with HCl) are mixed with 14 μL RNA (3-10 μg); mixed completely by vertexing, and then incubated for 45 minutes at 35°C to block. The samples were briefly allowed to cool to room temperature (2-3 min), then working quickly, 1 μL 30 mM aldehyde reactive probe (ARP/aminooxy biotin, 10009350, Cayman Chemicals, stock made in water) is added first, then 2μL of 7.5 mM NaIO_4_ (periodate, stock made in water) is added. The periodate is allowed to perform oxidation for exactly 10 minutes at room temperature in the dark. The periodate is then quenched by adding 3 μL of 22 mM sodium sulfite (stock made in water). The quenching reaction is allowed to proceed for 5 minutes at 25°C. Next, the reactions are moved back to the 35°C heat block, and the ligation reaction is allowed to occur for 90 min. The reaction is then cleaned up using a Zymo-I column. 26 μL of water is added in order to bring the reaction volume to 60 μL, and the Zymo protocol is followed as per the above details. The RNA is then eluted from the column using 20 μL water.

### LBD

The step-by-step protocol for LBD was summarized in extended data Fig. 4.

### Reagents

LEL and other lectins, Ac4ManNAZ, DPEG-Biotin were purchased from Vector Laboratories; Human tissue RNAs were purchased from Biochain Institute; Human thymus RNA, plant RNA and amniotic fluid were purchased from Amsbio. NorthernMax kit was from Invitrogen; Protein-free blocking buffer was from Li-Cor Company; m-PEG3-aldehyde was from Creative labs; Anti-biotin-HRP and NC membrane were from Thermo Fisher Scientific; All cell lines were from ATCC and provided by colleagues. HUVECs were from Lonza.

## ACKNOWLEDGMENTS

We thank Drs. Tony Wang, Daping Fan, Kun Chen, Jianming Qiu, Shizhen Wang, Cuthbert Simpkins, Nihar Nayak, Maria L. Spletter and Brain D. Ackley for kindly providing cell lines and/or RNA samples. This project was supported by National Institutes of Health (R15AI138116, to MF), UMKC Funding for Excellence (2024) and SPiRE (2024, to M.F).

## AUTHOR CONTRIBUTIONS

M.F. conceived, designed and supervised the overall study. Y.L and M.F performed most of the experiments. Y. Q performed metabolic labeling experiments for mouse. T.L and P.N help to perform mouse tissue expression experiments, and M.F. wrote the manuscript.

---

**Extended data Figure 1.**
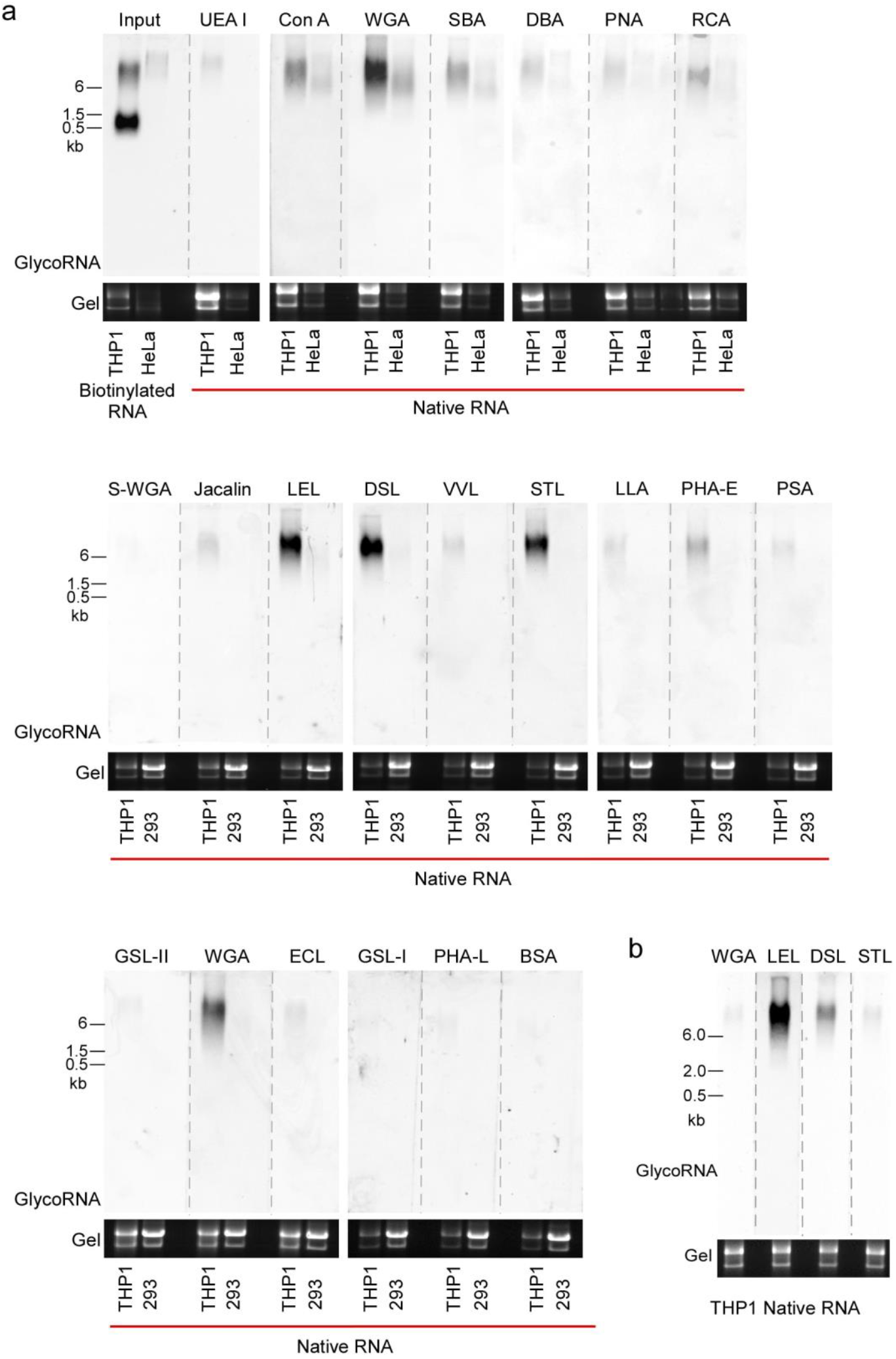
Screening lectins that can be used for detection of glycoRNAs. 20 different lectins were purchased from Vector Laboratories. Total RNAs isolated from THP1, HeLa or HEK293 cells were used for screening. Input RNAs isolated from Ac4ManNAz-labeled THP1 cells was conjugated with biotin and detected by anti-biotin-HRP. All lectin concentrations are 1.5 μg/ml.

**Extended data Figure 2.**
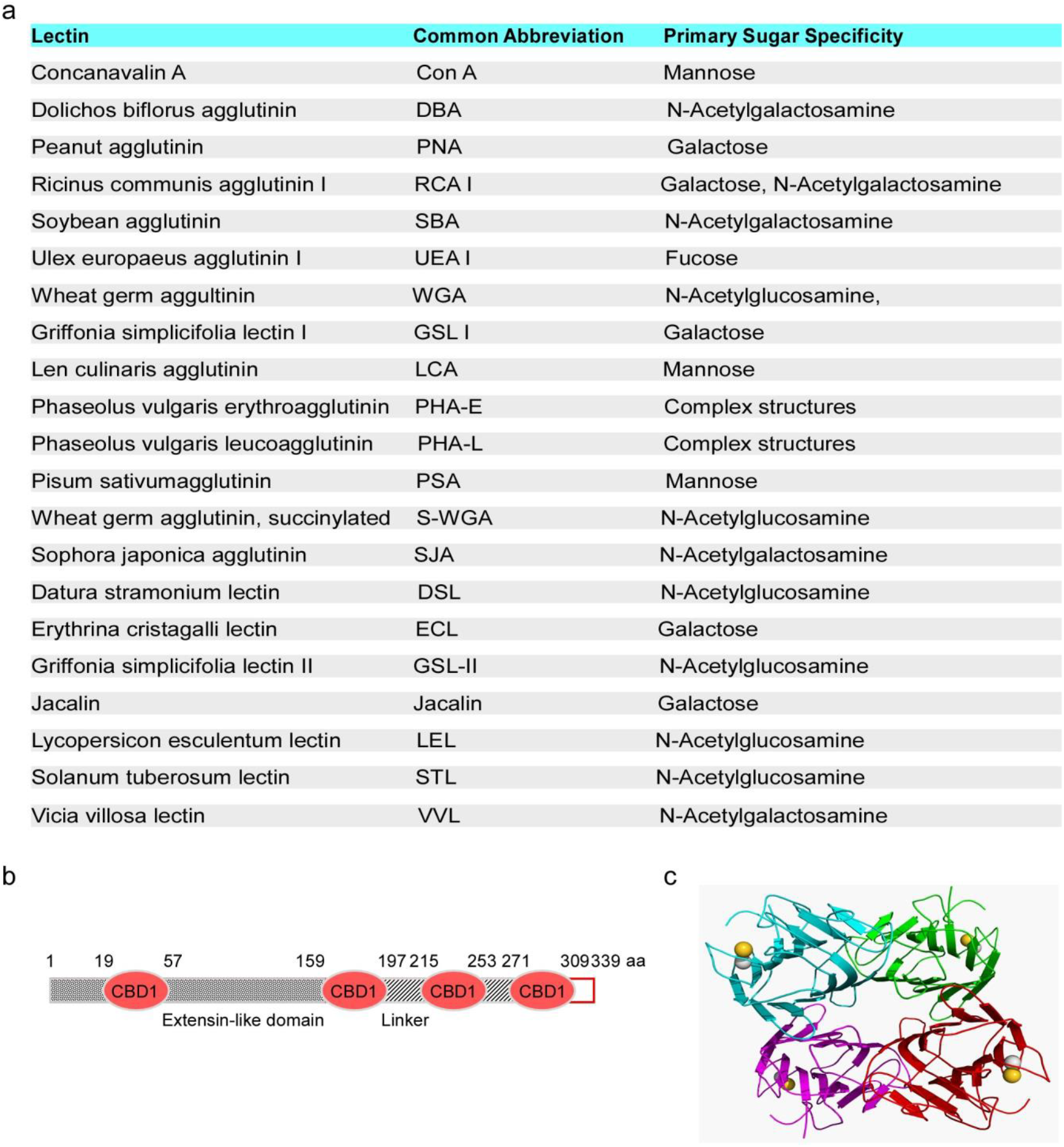
Information of lectins and LEL. (a) The full names and abbreviations and primary sugar specificity of 20 different lectins (Vector Laboratories). (b) The schematic of LEL domain structures. CBD1: chitin-binding domain. (c) The crystal structure of a lectin (adapted from Wikipedia).

**Extended data Figure 3.**
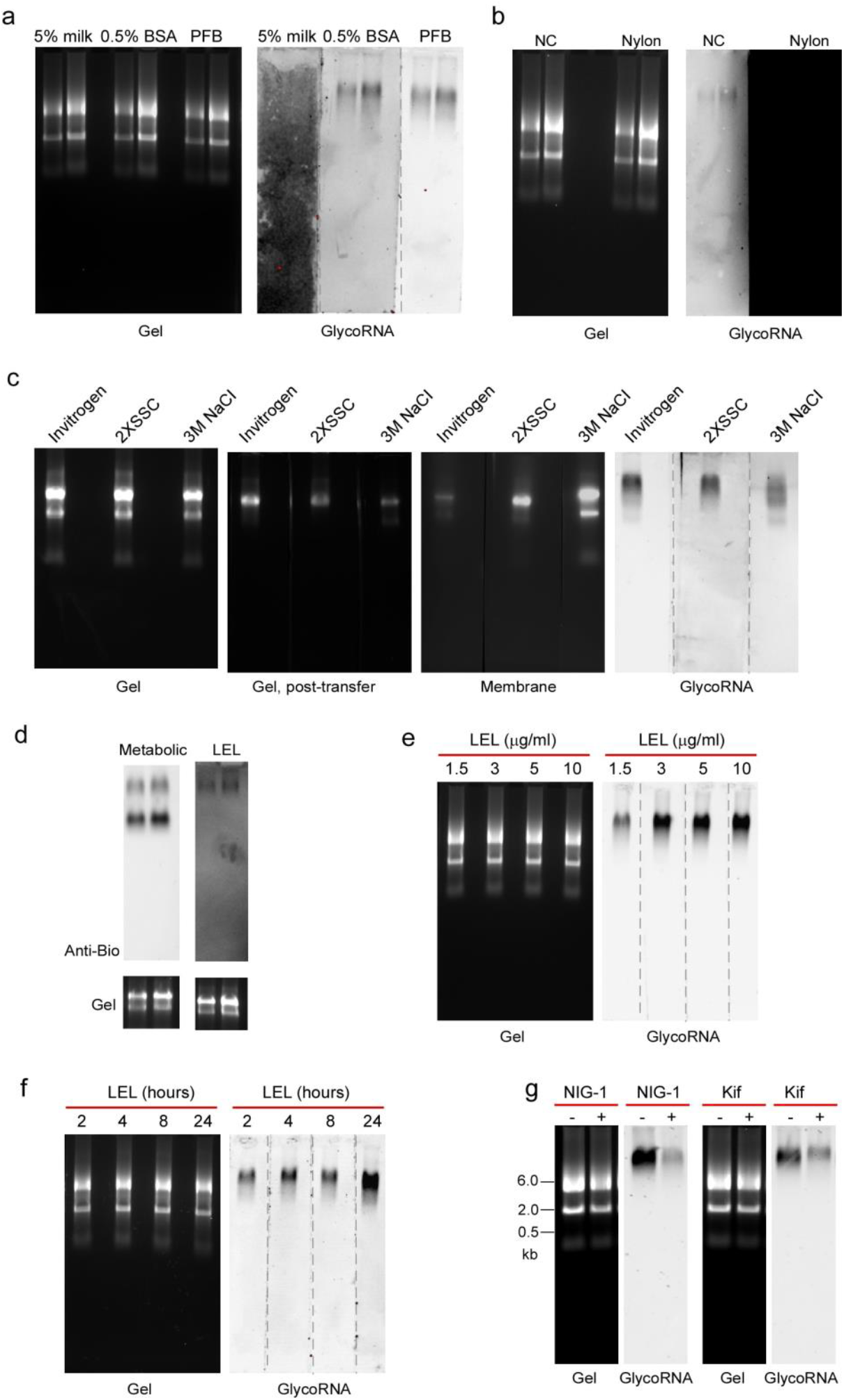
Optimizing the condition of lectin-based detection. (a) Three types of blocking buffers as indicated were compared. (b) Two types of membrane as indicated were compared. (c) Three types of transfer buffers as indicated were compared. (d) General agarose from Thermo Fisher Scientific was used for metabolic labeling analysis and LBD. (e) Different concentrations of LEL as indicated were compared. (f) Different incubation times of LEL (3 μg/ml) were compared. (g) Total RNAs isolated from THP1 cells treated with or without NIG-1 (4 μM/L) or Kifunensine (1 μM/L, Kif) were analyzed by LBD.

**Extended data Figure 4.**
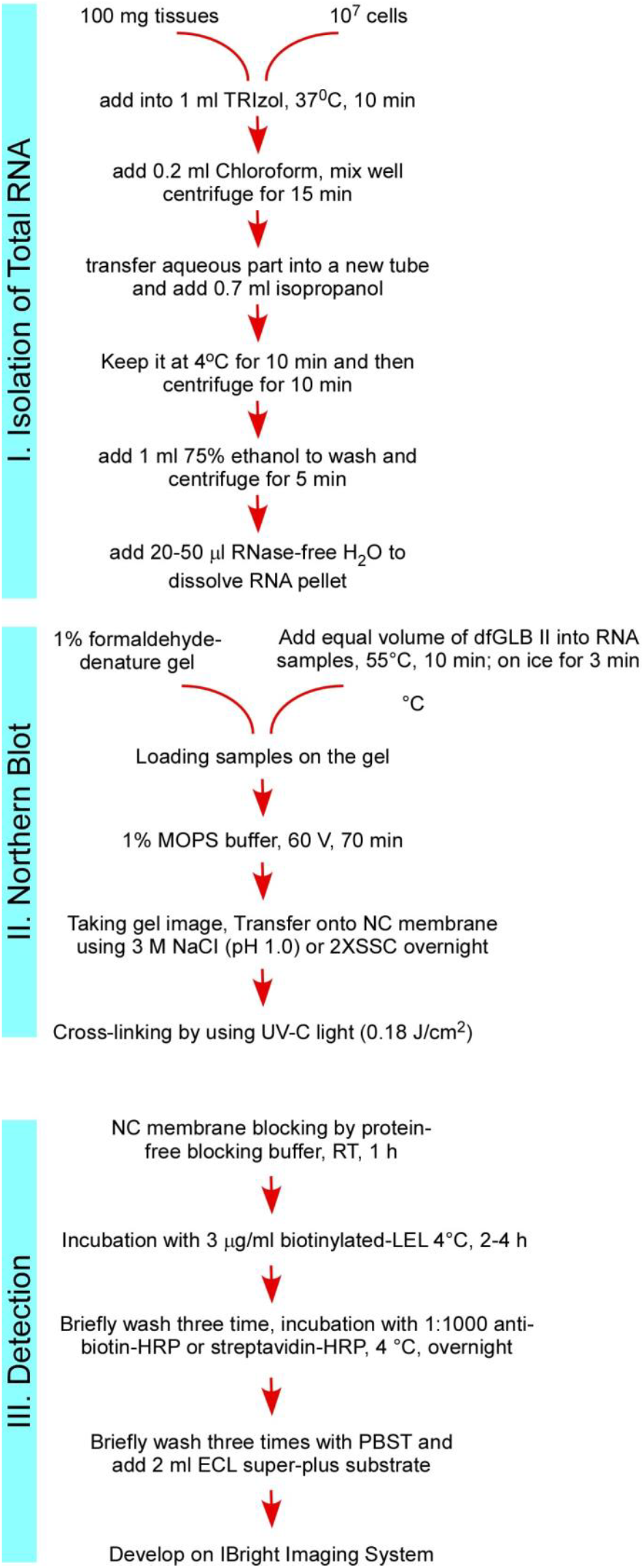
The step-by-step protocol for detection of native glycoRNAs by lectin-based method.

**Extended data Figure 5.**
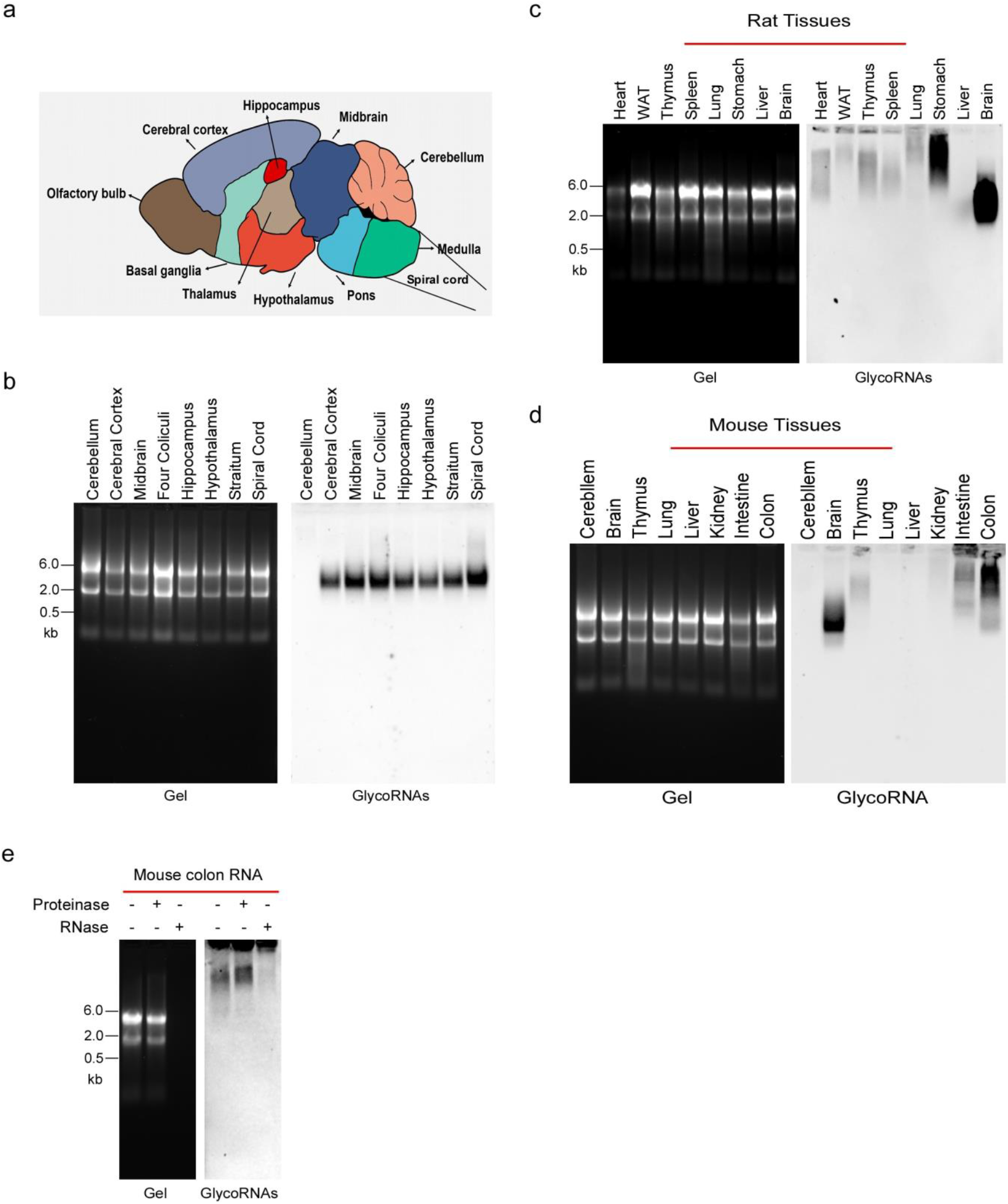
Expression of glycoRNAs in mouse and rat tissues. (a) Total RNAs were isolated from different regions of mouse brain. (b) The expression of glycoRNAs in the different regions of mouse brain were analyzed by LBD. (c) The expression of glycoRNAs in the different rat tissues were analyzed by LBD. (d) The expression of glycoRNAs in the different mouse tissues were analyzed by WGA. (e) Mouse colon RNAs were treated with or without Proteinase K or RNase and then analyzed by LBD.

**Extended data Figure 6.**
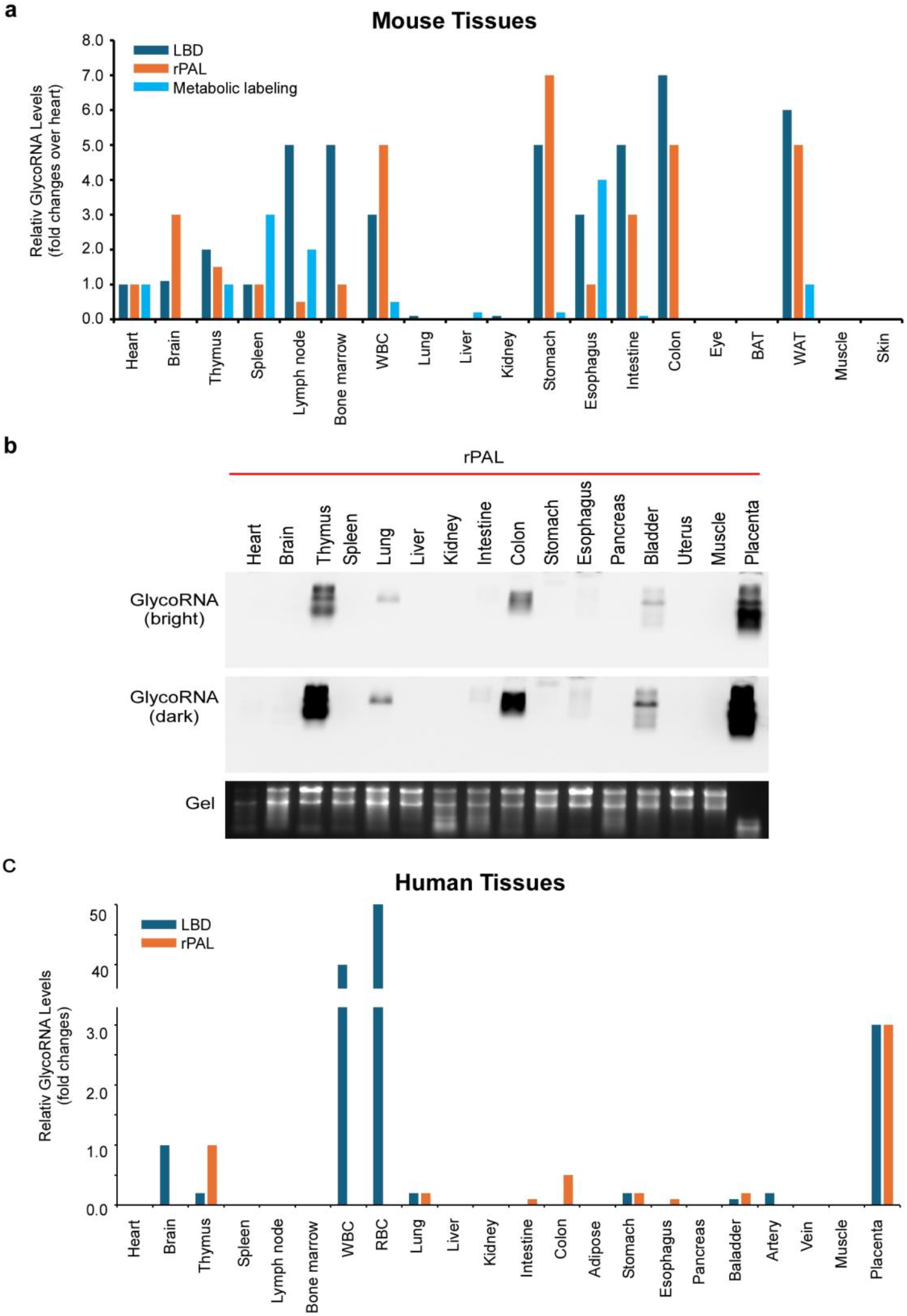
Expression patterns of glycoRNAs in mouse and human tissues. (a) The Quantification of glycoRNAs in mouse tissues were analyzed by three methods: LBD, rPAL and metabolic labeling as indicated. 10-15 μg total RNAs were used. The experiments were repeated at least twice. (b) The 16 human tissue RNAs were profiled by rPAL. (c) The quantification of glycoRNAs in human tissues were analyzed by LBD and rPAL. 8-10 μg total RNAs were used.

**Extended data Figure 7.**
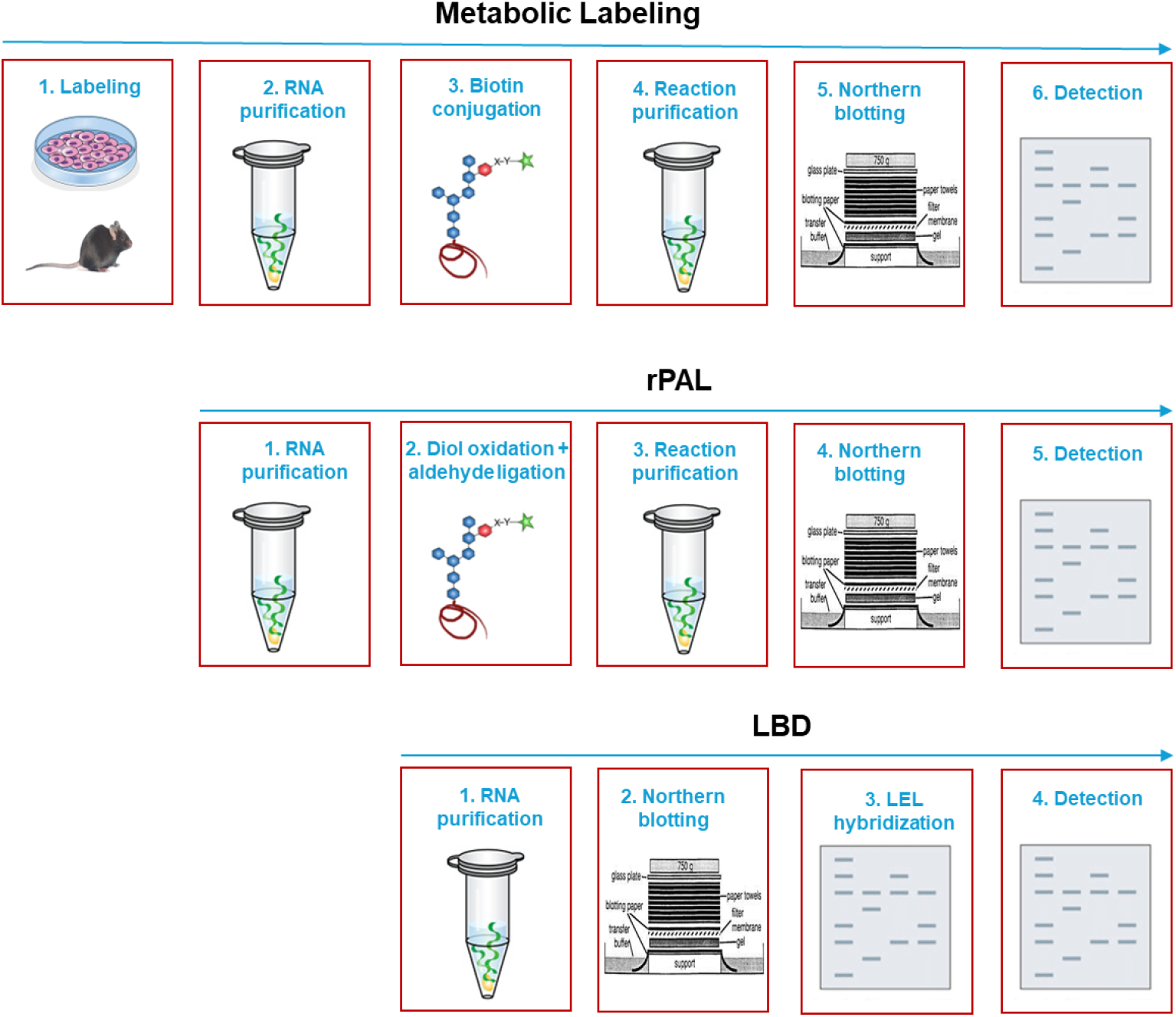
Schematic comparison of metabolic labeling, rPAL and LBD procedures.

